# Butyrate regulates the blood-brain barrier transport and intra-endothelial accumulation of Alzheimer’s disease Amyloid-beta peptides

**DOI:** 10.1101/2025.10.24.684335

**Authors:** Vaishnavi Veerareddy, Zengtao Wang, Purna C. Kashyap, Karunya K. Kandimalla

**Affiliations:** Department of Pharmaceutics and Brain Barriers Research Center, University of Minnesota, College of Pharmacy, Minneapolis, MN; Department of Gastroenterology and Hepatology Department of Medicine, Mayo Clinic, Rochester, Minnesota, MN

**Keywords:** Alzheimer’s, Amyloid beta, Blood-brain barrier, Butyrate, Insulin signaling, P-gp

## Abstract

Alzheimer’s disease (AD) is characterized by the pathological deposition of amyloid beta (Aꞵ) proteins as amyloid plaques, tau aggregates, and cerebrovascular dysfunction that drive disease progression. Butyrate, a gut microbial metabolite, has been found to be reduced in AD patients; butyrate supplementation improved cognition and decreased amyloid burden in animal models. However, the precise underlying mechanisms are unclear. Our previous studies have demonstrated that insulin signaling impacts Aꞵ transport kinetics at the blood-brain barrier (BBB). In this study, we investigated the effect of butyrate treatment on intra-endothelial Aꞵ accumulation and BBB integrity by modulating the insulin signaling pathway. The effect of butyrate on Aꞵ accumulation was assessed by flow cytometry in BBB cell culture models. Insulin signaling activation and the expression of various receptors and transporters at the BBB were evaluated by Western blots and confocal microscopy. The roles of various molecular mediators were confirmed using specific inhibitors (MK2206, Trametinib, Rapamycin, VX-745). The effect of butyrate on the expression of BBB receptors and transporters that play a critical role in Aꞵ trafficking was examined in mouse brains colonized with butyrate-producing bacteria via immunohistochemistry. Butyrate significantly decreased Aβ42 accumulation in endothelial cells. This effect was associated with insulin signaling pathway activation, particularly AKT and ERK phosphorylation. Inhibitor studies established the critical role of these specific arms, as co-incubation with MK2206 (AKT inhibitor) or Trametinib (ERK inhibitor) reversed the protective effect of butyrate and increased Aβ42 accumulation. However, mTOR and p38 inhibitors did not show a similar effect. In addition, butyrate restored P-glycoprotein efflux transporter expression and claudin-5 tight junction protein levels that were reduced with Aβ treatment. These effects were supported by in vivo work, which demonstrated the upregulation of Tissue Inhibitor of Metalloproteinases-2 (TIMP-2). This protein is associated with AKT activation and extracellular matrix stabilization in mice colonized with butyrate-producing bacteria. In conclusion, we have demonstrated that butyrate decreases Aβ42 uptake at the BBB endothelium by activating the AKT and ERK arms of the insulin signaling pathway. These changes may also improve the integrity of BBB tight junctions by increasing claudin-5 expression and extracellular matrix, and by upregulating TIMP-2 expression. This study highlights butyrate’s potential as a therapeutic modulator of AD-related BBB dysfunction.

## Introduction

Alzheimer’s disease (AD) is an irreversible, progressive neurodegenerative disorder and the leading cause of dementia in the elderly^1^. The major hallmarks of AD include amyloid plaques in brain parenchyma, cerebral amyloid angiopathy, characterized by vascular amyloid beta (Aβ) deposits, and hyperphosphorylated tau tangles in neurons^2^. Key risk factors such as age, cerebrovascular dysfunction, insulin resistance, and gut dysbiosis are known to worsen these biomarkers, especially amyloid burden^3^. While overproduction of Aβ peptides primarily drives familial AD, amyloidosis in sporadic AD is attributed mainly to impaired Aβ clearance^2^.

The blood-brain barrier (BBB) plays a dynamic role in maintaining brain homeostasis. Clearance of Aβ peptides from the brain is mediated by the low-density lipoprotein receptor-related protein 1 (LRP1) and Permeability-glycoprotein (P-gp), whereas plasma-to-brain Aβ influx is mediated by receptor for advanced glycation end products (RAGE) in the BBB endothelium^4,5^. In the AD brain, this homeostasis is disrupted, and experimentally evident by reduced LRP1 and P-gp expression along with elevated RAGE levels, leading to increased Aβ deposition in the brain and most likely in the BBB endothelium^6–8^. These Aβ deposits further compromise BBB integrity by increasing inflammatory cytokines^9^ and decreasing the tight junction proteins such as claudin-5 and occludin in endothelial cells^10^. Notably, the insulin signaling pathway, which regulates the function of these crucial transporters and tight junction expression, is also impaired by Aβ peptides^11,12^.

Although Aβ-targeting drugs like lecanemab and donanemab reduce amyloid burden and slow cognitive decline^13,14^, their effectiveness in restoring BBB integrity and modulating the complex AD pathology remains incomplete. This highlights the urgent need to identify novel approaches that naturally restore BBB function, that complement existing treatments.

The gut microbial metabolite butyrate is emerging as a compelling therapeutic candidate due to its neuroprotective effect via the gut-brain axis. Produced by Clostridium bacterial clusters via the anaerobic fermentation of dietary fiber^15^, butyrate production is significantly reduced in AD patients, which is correlated with cognitive decline. As a histone deacetylase (HDAC) inhibitor, butyrate was shown to improve cognitive function in transgenic AD mouse models^16,17^, protect against Aβ toxicity, and reduce neuroinflammation ^18,19^. Further, the butyrate effect was verified in animal models, where leaky brain vasculature, typically observed in germ-free mice, is repaired when a butyrate-producing microbiome colonizes the mice^20^. However, this process is disrupted due to dysbiosis in AD patients, where the presence of the butyrate-producing microbiome is significantly reduced, which was positively correlated with cognitive decline and disease progression scores^21^. Notably, butyrate repairs BBB integrity, increasing the expression of tight junction proteins-claudin-5 and zona occludens-1 in vitro^22,23^, and reversing the leaky vasculature observed in germ-free mice in vivo^20^. Strategies to increase gastrointestinal absorption of butyrate into systemic circulation were shown to improve neuroinflammation and ameliorate brain ischemic injury^24,25^. Apart from its central nervous system benefits, butyrate was known to increase peripheral insulin sensitivity by activating the insulin signaling pathway^26^. Apart from these protective effects, conflicting results exist, which could be due to the complexity of AD pathology. Therefore, it is crucial to elucidate the specific molecular connection between butyrate, insulin signaling, and Aβ trafficking at the BBB, which has not been established.

In this study, we investigated the precise mechanism of action of butyrate in modulating toxic Aβ accumulation in polarized BBB endothelial cell culture models. Furthermore, we employ germ-free mice colonized with butyrate-producing or butyrate-deficient microbiome to investigate the expression changes of key proteins in the brain microvasculature, thus providing a comprehensive view of butyrate’s impact on BBB physiology.

## Methods

### Cell culture

The immortalized human cerebral microvascular endothelial cells (hCMEC/D3) were gifted by P-O Couraud (Institute Cochin, France) and were cultured as previously reported^27^. The endothelial cell basal medium (Cell Applications, CA) supplemented with 1% v/v lipid concentrate (Thermo Fisher Scientific, MA), 1% v/v penicillin-streptomycin (Sigma-Aldrich, St. Louis, MO), 1 ng/mL recombinant human fibroblast growth factor-basic (PeproTech, Rocky Hill, NJ), 1.4 μM hydrocortisone reconstituted in ethanol (Sigma-Aldrich, MO), 5 μg/mL ascorbic acid (Sigma-Aldrich, MO), 10 mM HEPES (Millipore Sigma, MO), and 5% fetal bovine serum (FBS) (Atlanta Biologicals, GA), referred as D3 medium was used to culture the cells. The polarized hCMEC/D3 cell monolayers (passage 35) were grown in coverslip-bottomed dishes and cell culture plates coated with collagen (5 μg/ Cm^2^) (Thermo Fisher Scientific, Waltham, MA). Cells were starved in 1% serum-containing D3 medium for 14-16 h before all experiments to reduce background activation of signaling pathways.

Primary bovine brain microvascular endothelial (BBME) cells were acquired from Cell Applications Inc. (San Diego, CA). The cells were cultured on 6-well plates coated with collagen (5 μg/ Cm^2^) (Thermo Fisher Scientific, MA) followed by 0.01% bovine fibronectin until >90% confluency is attained. The BBME cells were grown in Dulbecco’s Modified Eagle’s medium (DMEM)/F12 medium (Cellgro, Corning, NY) supplemented with donor horse serum (Cellgro, Corning, NY), gentamycin (25 mg) (Cellgro, Corning, NY), sigma heparin sodium salt (50 mg) (Sigma-Aldrich, St. Louis, MO).

### Aβ40/42 solution preparation

The Fluorescein Isothiocyanate (F)-labeled Aβ40, unlabeled Aβ40, F-Aꞵ42, and unlabeled Aꞵ42 were supplied by Aapptec (Louisville, KY). Monomers were prepared by dissolving them in 1,1,1,3,3,3, - hexafluoro-2-propanol (HFIP) (Sigma-Aldrich, MO) as reported previously^28^. A clear peptide solution was aliquoted into glass vials and dried overnight in the fume hood. The film thus formed was dried under vacuum and stored at -20 °C over desiccant. The Aβ film was dissolved in dimethyl sulfoxide, alkaline water, and F-12 (R&D systems, MN) before the experiment. Later, the Aβ solution was centrifuged for 5 min at 6000 rpm to eliminate high molecular species^29^. The supernatant was then reconstituted in DMEM (Gibco, NY) and supplemented for the cells. In the cell culture experiments, precautions were taken to limit dimethyl sulfoxide exposure from Aβ solutions to less than 0.25%.

### Flow cytometry

#### Butyrate effect on F-Aβ40 or F-Aβ42 accumulation in BBB endothelium in vitro

The polarized hCMEC/D3 cell monolayers cultured in 6-well plates were treated with butyrate, 10 nM (sodium butyrate, Sigma-Aldrich, MO) for 1 h, 5 h, and 23 h in low-serum D3 medium, followed by F-Aβ40 (2 μM) or F-Aβ42 (1 μM) treatment for one hour. BBME cells cultured in 6-well plates were treated with butyrate (10 nM) for 5.5 h in low-serum D3 medium, followed by F-Aβ42 (1 μM) co-incubation for 30 minutes. After treatment, cells were washed with chilled PBS twice. Later, they were trypsinized for 3 min at 37 °C and quenched with FBS. Then the cell suspension was diluted with chilled PBS and centrifuged for 5 min at 1200 rpm. Following that, they were resuspended in PBS and fixed with 4% paraformaldehyde (PFA)^30^. Cells were gently vortexed and fluorescence accumulation was acquired with a Flow cytometer (Laser source: 100 mW Blue, 488 nm, Excitation: 495 nm, Emission: 519 nm with 530/30 filter) and analyzed in FlowJo v10.0. The F-Aβ40 or F-Aβ42 uptake was represented as histograms of intracellular fluorescent intensities normalized to the mode. Fold change was calculated by dividing the individual group by the untreated control, F-Aβ group.

#### Effect of insulin signaling mediators on F-Aβ42 accumulation in polarized hCMEC/D3 Monolayers

The polarized hCMEC/D3 monolayers cultured in 6-well plates were incubated for 5.5 h under the following conditions in low-serum D3 medium: control (no treatment), butyrate alone (10 nM), butyrate + inhibitor, or inhibitor alone. Following that, the cells were incubated in colorless DMEM with the same treatment as preceding 5.5 hours, along with F-Aβ42 (1 μM) and insulin (50 nM) for 30 minutes. A separate experiment was conducted for each of these kinase phosphorylation inhibitors: 10 μM MK2206 (AKT), 100 nM Rapamycin (mTOR), 0.5 μM trametinib (MEK), 10 nM VX-745 (p38), obtained from Selleckchem, TX. Samples for flow cytometry were processed and investigated as mentioned earlier.

### Membrane fractionation

The polarized hCMEC/D3 cell monolayers were grown in 150 mm culture dishes (CELLTREAT scientific products, MA) containing D3 media until they were fully confluent. Cells were treated with butyrate in low serum D3 medium for 5 h, followed by butyrate and Aβ42 (1 μM) co-incubation in DMEM for 1 h. After treatment, cells were washed 3 times with ice-cold PBS. Plates were scraped with PBS (1 ml/*3) and lysates were collected into a 15 ml falcon tube. The resultant cell suspension was centrifuged for 5 min at 1200 rpm (Thermo Scientific IEC CL40). Later, the supernatant was aspirated, and the pellet was processed for membrane protein isolation (Minute™ Plasma Membrane Protein Isolation and Cell Fractionation Kit, Invent Biotechnologies, MN). The pellet was resuspended in the lysis buffer supplemented with RIPA buffer, Protease Inhibitor Cocktail 100x (Sigma-Aldrich, MO), phosphatase inhibitor cocktail 100x (Santa Cruz, TX), and Pierce™ nuclease 100x (Thermo Fisher Scientific, MA). The cell suspension was filtered through a cartridge and vortexed to separate nuclei. The supernatant was collected and centrifuged for 1 h at 16000*g at 4 °C to separate the cytosol fraction (supernatant) from the total membrane (pellet). The pellet was dissolved in a Minute denaturing protein solubilization reagent along with a protease and phosphatase inhibitor cocktail. The protein concentration of the total membrane fraction was determined using a Pierce^TM^ bicinchoninic acid assay kit (Thermo Fisher Scientific, MA). About 10-20 µg of membrane fraction protein was loaded on the gel for the western blot.

### Western blot

The hCMEC/D3 cells grown in 6-well plates were treated with butyrate (10 nM) alone for 6 h or butyrate with insulin (50 nM) stimulation for the last 5 minutes or insulin alone in low serum D3 medium. Following the experiment, cells were washed with ice-cold PBS, and lysates were collected using lysis buffer, and protein concentration was measured. Whole-cell lysates (25 µg) and total membrane protein (20 µg) were loaded on 4–12% Criterion™ XT Bis-Tris protein gel (Bio-Rad Laboratories, CA) with 20x, XT MOPS running buffer (Bio-Rad, Hercules, CA) for 1.5 h. Later, the gel was transferred to nitrocellulose membrane at 100 V for 30 min using methanol, deionized water, and tris/glycine buffer (Bio-Rad, Hercules, CA) in the ratio of 2:7:1. The membrane was blocked for one hour with 5% v/v blotting grade blocker (Bio-Rad, CA) in tris buffered saline (TBS) containing Tween20 (0.1%) (TBST) (Bio-Rad, CA). Then, the membrane was washed with TBST and was incubated overnight at 4 °C with one of the primary antibodies P-AKT/Ser473 (#4060), MDR1/ABCB1/P-glycoprotein (#13978), Phospho-ERK1/2 (#4377), GAPDH (#5174) from Cell Signaling Technology, Danvers, MA, or receptor for advanced glycation end products (RAGE) (#AB3611, Abcam, Cambridge, MA)] diluted with 5% v/v BSA (Sigma-Aldrich, St. Louis, MO). Antibodies were diluted with 5% BSA in TBST at 1:1000 dilution. The following day, the membrane was washed with TBST and incubated for 1 h at room temperature with a secondary antibody conjugated with near-infrared (800 nm) (Licor; Lincoln, NE) at 1:2000 dilution. Following incubation, the membrane was washed with TBST and TBS and imaged using an Odyssey Licor imager. Protein bands were quantified using Image Studio Lite 5.2. and normalized using loading protein concentrations of GAPDH for whole cell lysates and calnexin for membrane fraction.

### Immunocytochemistry

The hCMEC/D3 cells monolayers were cultured on 35 mm round coverslip bottom dishes in D3 medium containing 5% FBS. A day before the experiment, cells were transitioned to 1% FBS-containing D3 medium. Cells were pre-treated with either blank or butyrate (10 nM) in low serum D3 medium for 5 h. Following pre-treatment, the medium was replaced with colorless DMEM. For the experimental groups, butyrate was re-administered, and cells were co-incubated with Aβ42 (1 μM) for the butyrate group or treated with Aβ42 (1 μM) alone for the control group for an hour. Following treatment, cells were washed with ice-cold PBS and fixed with 4% PFA and blocked using 10% goat serum for an hour. Following that, the cells were incubated overnight at 4 °C with the primary antibody, claudin-5 (1:50, #49564, Cell Signaling Technology, Danvers, MA). The next day, the cells were washed and incubated with Rabbit anti-Goat IgG AF647 secondary antibody (1:1000, #A-21446 Invitrogen, Waltham, MA) for an hour in the dark. Later, they were stained with DAPI (2 µM, Invitrogen, Waltham, MA) for 10 minutes and washed. They were mounted using Prolong Gold antifade mounting medium (Invitrogen, Waltham, MA). Imaging was performed at the University of Minnesota Imaging Center in Nikon Confocal A1Rsi NSIM confocal microscope using a 60X,1.4NA objective.

### Animal models

Germ-free animals, BL6 mice, were grown in an aseptic facility in Mayo in accordance with IACUC#A00005635^31^. They were colonized with either butyrate-producing Clostridium senegalense DSM 25507 WT or double-knockout (KamD and crotonase genes) mutant incapable of biosynthesizing butyrate by oral gavage culture at 3.5-4.5 weeks of age. They were maintained for 2 weeks in the same facility before harvesting brains.

### Immunohistochemistry (IHC)

The mice’s brains were formalin-fixed and embedded in paraffin, and then sectioned into 5-µm-thick slices on a slide. Sections were processed for IHC as previously published^32^. Brain tissue was stained for tissue inhibitors of metalloproteinases (TIMP-2, 1:100 dilution, #5738 from Cell Signaling Technology, MA) with 3,3′-Diaminobenzidine (DAB) chromogen. Images were acquired using a Zeiss Axio Scan.Z1 microscope at 20X magnification. Images were analyzed in QuPath-0.5.1 where butyrate KO mouse brains were compared to butyrate-producing microbiome colonized mouse brains (n=4). Mean DAB pixel intensity from both groups was obtained by averaging over 100 cell-specific regions of interest in the brain cortex, followed by pair-wise comparison.

### Statistical analysis

All statistical analyses were performed using GraphPad Prism software (version 10.0). Multiple groups were compared using one-way ANOVA followed by Bonferroni post-test, and the differences in the means of the two groups were compared using Student’s T-test. IHC data were compared by paired t-test. The level of significance is indicated as follows: *p-value<0.05, **p-value<0.01, ***p<0.001.

## Results

### 1. Butyrate effect on F-Aβ40 or F-Aβ42 accumulation in BBB endothelium in vitro

The polarized hCMEC/D3 cell monolayers were pre-treated with butyrate for various time periods to investigate its impact on F-Aβ40 or F-Aβ42 uptake. Fold changes of cellular median fluorescence intensity (MFI) from three replicates in each group were compared. The 6-h butyrate treatment significantly decreased F-Aβ40 accumulation (0.89-fold, p < 0.05), whereas 2-h and 24-h pretreatments had no significant effect compared to the control group (Figure 1A & B). On the other hand, butyrate significantly decreased F-Aβ42 accumulation at both 2-h (0.5-fold, p < 0.01) and 6-h time points (0.5-fold, p < 0.01), with no effect observed at the 24-h treatment (Figure 1C & D). As butyrate was effective in reducing both F-Aβ40 and F-Aβ42 in hCMEC/D3 cell monolayers after 6 h of treatment, its effect was further verified in a primary BBB cell culture model constructed from BBME cells. These studies have shown that butyrate significantly reduced F-Aβ42 accumulation to 0.6-fold (p<0.01) compared to the control group (Figure 1E, F).

**Figure 1:**
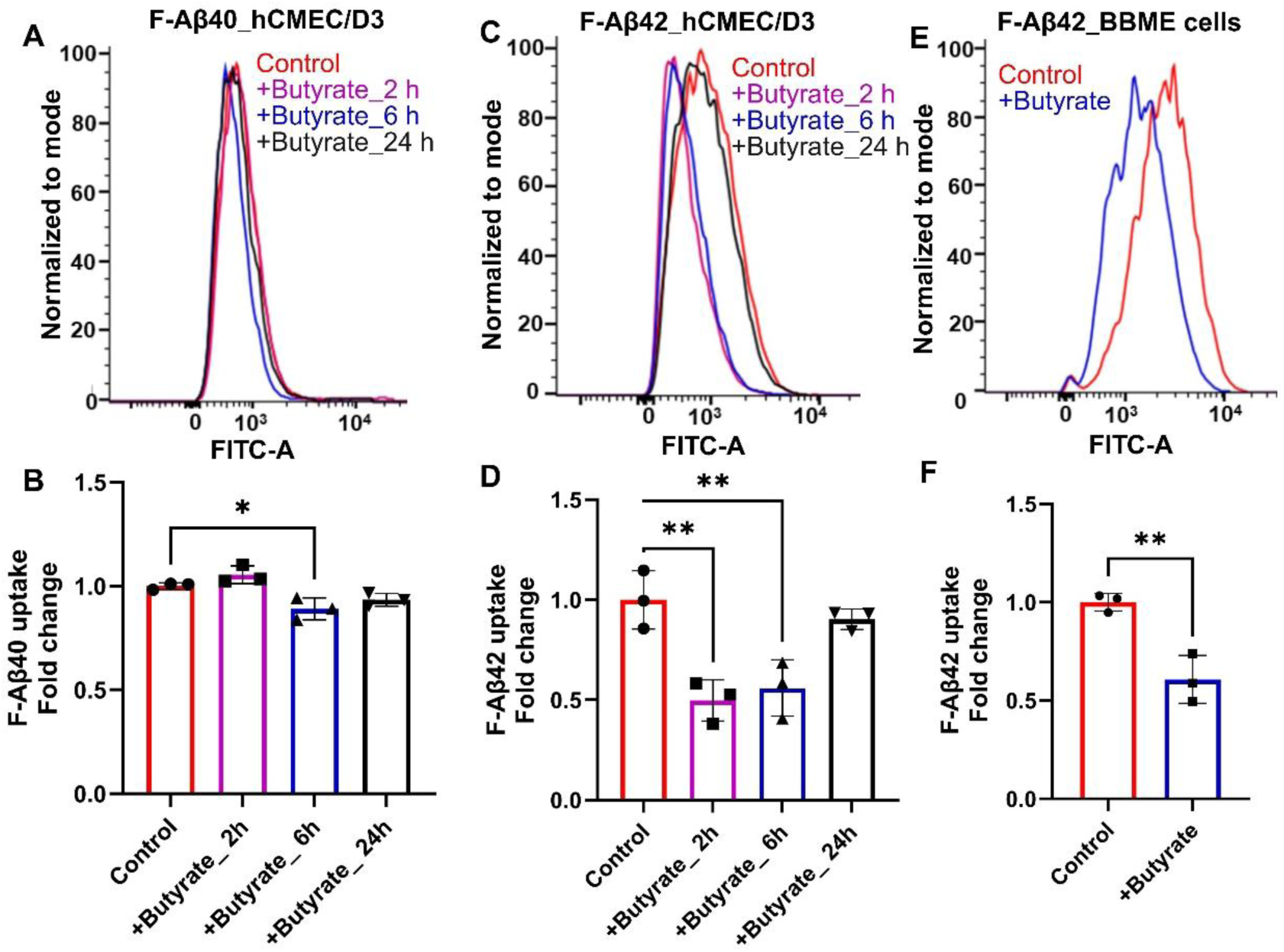
F-Aβ40 or Aβ42 accumulation was reduced in the BBB endothelial cell culture models in vitro. The polarized hCMEC/D3 cell monolayers were pre-treated with 10 nM butyrate for 2 h, 6 h, and 24 h, with F-Aβ40 (2 μM) or F-Aβ42 (1 μM) added in the last hour of incubation. The BBME cells were treated with butyrate for 6 h, followed by co-incubation with F-Aβ42 (1 μM) for the last 30 minutes. Histograms represent the MFI of F-Aβ40 or F-Aβ42. Butyrate decreased F-Aβ40 accumulation at 6 h pre-treatment, (A & B) and F-Aβ42 accumulation at both 2 h and 6 h (C & D) in hCMEC/D3 cells. Butyrate decreased F-Aβ42 accumulation at 6 h in another BBB cell culture model constructed from BBME cells (E & F). Fold change of median fluorescence ± standard deviation was shown in bar charts. For comparing multiple groups, a One-way ANOVA followed by the Bonferroni multiple-comparison test was used, and the Student’s t-test was used for comparing two groups. *p-value<0.05, **p-value<0.01.

### 2. Effect of butyrate on AKT and ERK phosphorylation

The hCMEC/D3 cell monolayers were pretreated with butyrate for 6 h and then co-incubated with 50 nM insulin for 5 min. Insulin treatment increased AKT phosphorylation compared with the untreated group; however, this difference was not statistically significant. Co-incubation with insulin and butyrate significantly increased the AKT phosphorylation by 3-fold (p<0.05) in hCMEC/D3 monolayers compared to the insulin alone-treated group (Figure 2A & B). Moreover, insulin treatment significantly increased ERK phosphorylation by 3-fold (p<0.001) compared with the untreated group. The ERK phosphorylation was further increased upon incubation with butyrate and insulin (∼1.3-fold, p<0.05) compared to the insulin alone-treated group (Figure 2A & C).

**Figure 2:**
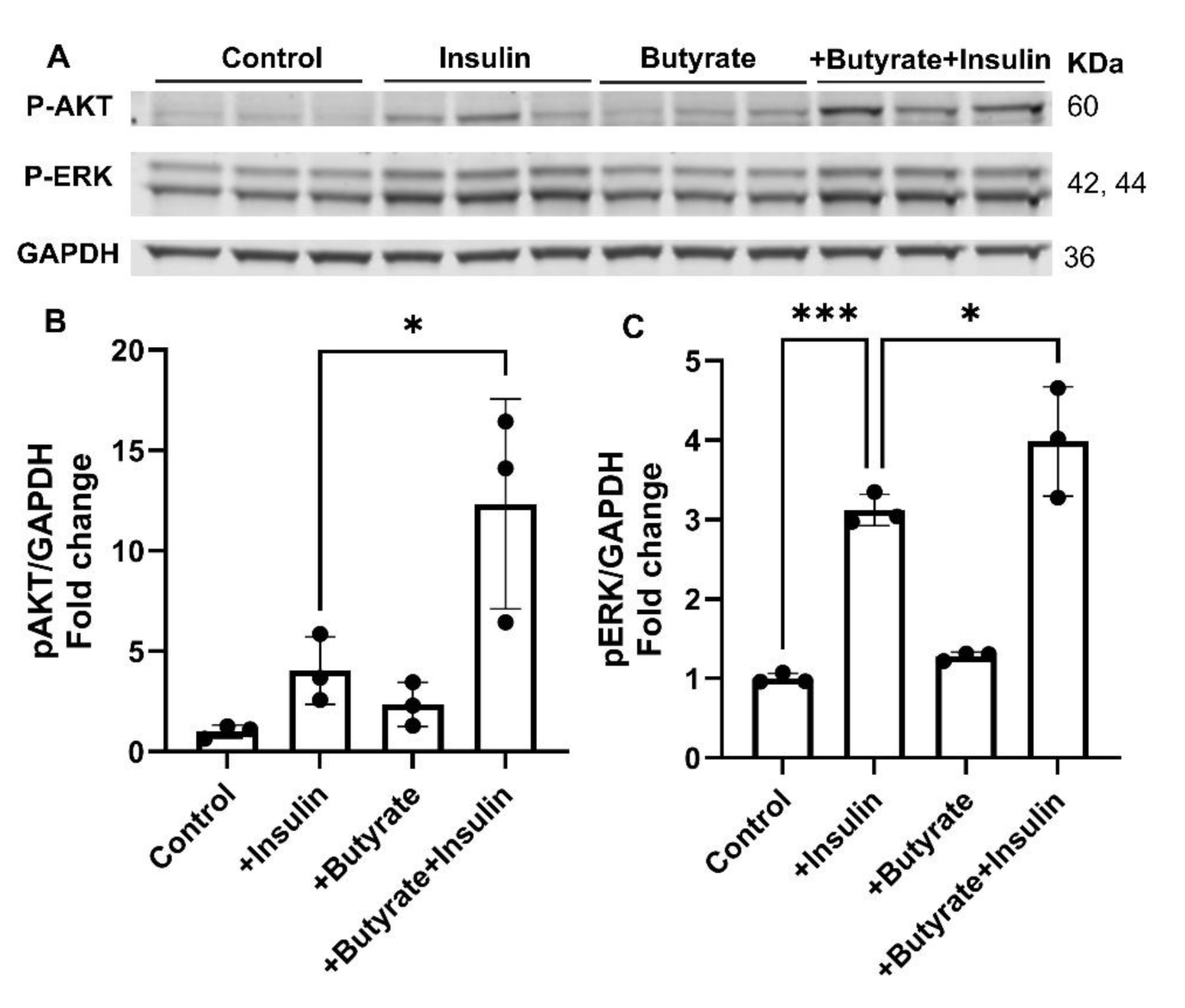
Butyrate augmented AKT and ERK phosphorylation stimulated by insulin in polarized hCMEC/D3 monolayers: Cells were treated with 10 nM butyrate for 6 h with or without 50 nM insulin stimulation for 5 min. A. Representative immunoblots showed increased AKT and ERK phosphorylation with 6-h butyrate pre-treatment and 5 min insulin stimulation compared to the group that received insulin treatment only. B & C. Quantification of phospho-AKT and ERK blots normalized to loading control, GAPDH, and presented as fold change. Mean ± standard deviation; One-way ANOVA followed by Bonferroni post-test; *p-value<0.05, **p-value<0.01.

### 3. Effect of insulin signaling mediators on F-Aβ42 accumulation in polarized hCMEC/D3 monolayers

To identify the role of molecular mediators activated by butyrate on intra-endothelial Aβ accumulation, the hCMEC/D3 cell monolayers were pre-incubated with butyrate and inhibitors of insulin signaling (AKT, mTOR, ERK) or stress signaling (p38) pathways for 6 h. Then they were co-incubated with insulin and F-Aβ42 for the last 30 minutes.

#### 3.1. The PI3K/AKT Pathway and F-Aβ42 Uptake

Upon butyrate treatment, the hCMEC/D3 cell monolayers demonstrated significantly lower F-Aβ42 accumulation (0.5-fold, p<0.05) compared to the untreated control. The F-Aβ42 accumulation increased in cell monolayers co-incubated with butyrate and MK2206 (AKT inhibitor) by 1.22-fold (p<0.05) compared to those treated with butyrate alone, which decreased F-Aβ42 uptake to 0.5-fold (Figure 3A & B). However, rapamycin (mTOR inhibitor) did not show any effect on F-Aβ42 accumulation in the hCMEC/D3 monolayers, as no significant difference in F-Aβ42 uptake was observed between the monolayers treated with butyrate alone and in combination with rapamycin (Figure 3C & D).

**Figure 3:**
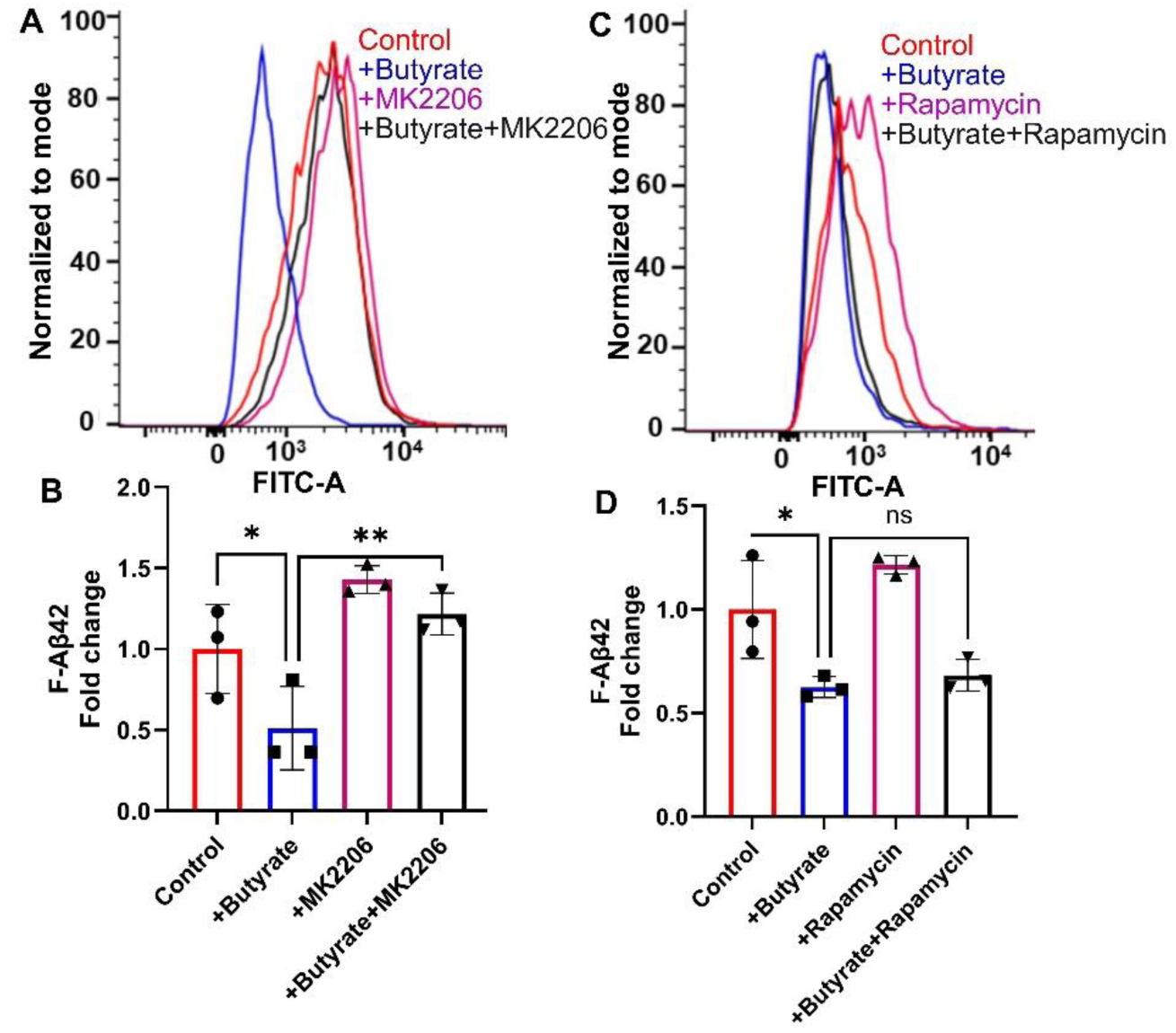
Effect of AKT and mTOR inhibitors on butyrate-mediated reduction of F-Aβ42 uptake in polarized hCMEC/D3 monolayers: Cells were incubated for 6 h under the following conditions: control (no treatment), butyrate alone (10 nM), butyrate + inhibitor, or inhibitor alone. In the last 30 min of each treatment, all groups received F-Aβ42 and insulin (50 nM). Histograms represent the MFI associated with F-Aβ42 uptake. Butyrate decreased F-Aβ42 accumulation compared to the control. A&B. Butyrate co-incubation with 10 µM MK2206 (AKT inhibitor) increased F-Aβ42 accumulation compared to butyrate alone, as shown in (A) histogram of intracellular fluorescence and (B) bar chart showing fold-change of median fluorescence ± standard deviation compared to the control. C & D. In another experiment, butyrate decreased F-Aβ42 accumulation compared to control, and co-incubation with 100 nM Rapamycin (mTOR inhibitor) did not alter F-Aβ42 accumulation, as shown in C histogram of intracellular fluorescence and D. Bar charts represent fold change of median fluorescence ± standard deviation. One-way ANOVA followed by Bonferroni post-test was performed; *p-value<0.05, **p-value<0.01.

#### 3.2. The ERK/MAPK pathway and F-Aβ42 uptake

Butyrate treatment significantly reduced F-Aβ42 accumulation (0.4-fold, p<0.05) in hCMEC/D3 monolayers compared to the control. In contrast, when cell monolayers are co-incubated with butyrate and trametinib, the F-Aβ42 uptake was the same as the control and is significantly higher (p<0.01) than butyrate alone treatment (Figure 4 A & B). We further investigated the role of another MAPK kinase, p38, which is activated predominantly under stress. We found that co-incubation of hCMEC/D3 monolayers with p38 inhibitor, VX-745, did not significantly affect F-Aβ42 accumulation compared to the butyrate treatment alone (Figure 4 C& D).

**Figure 4:**
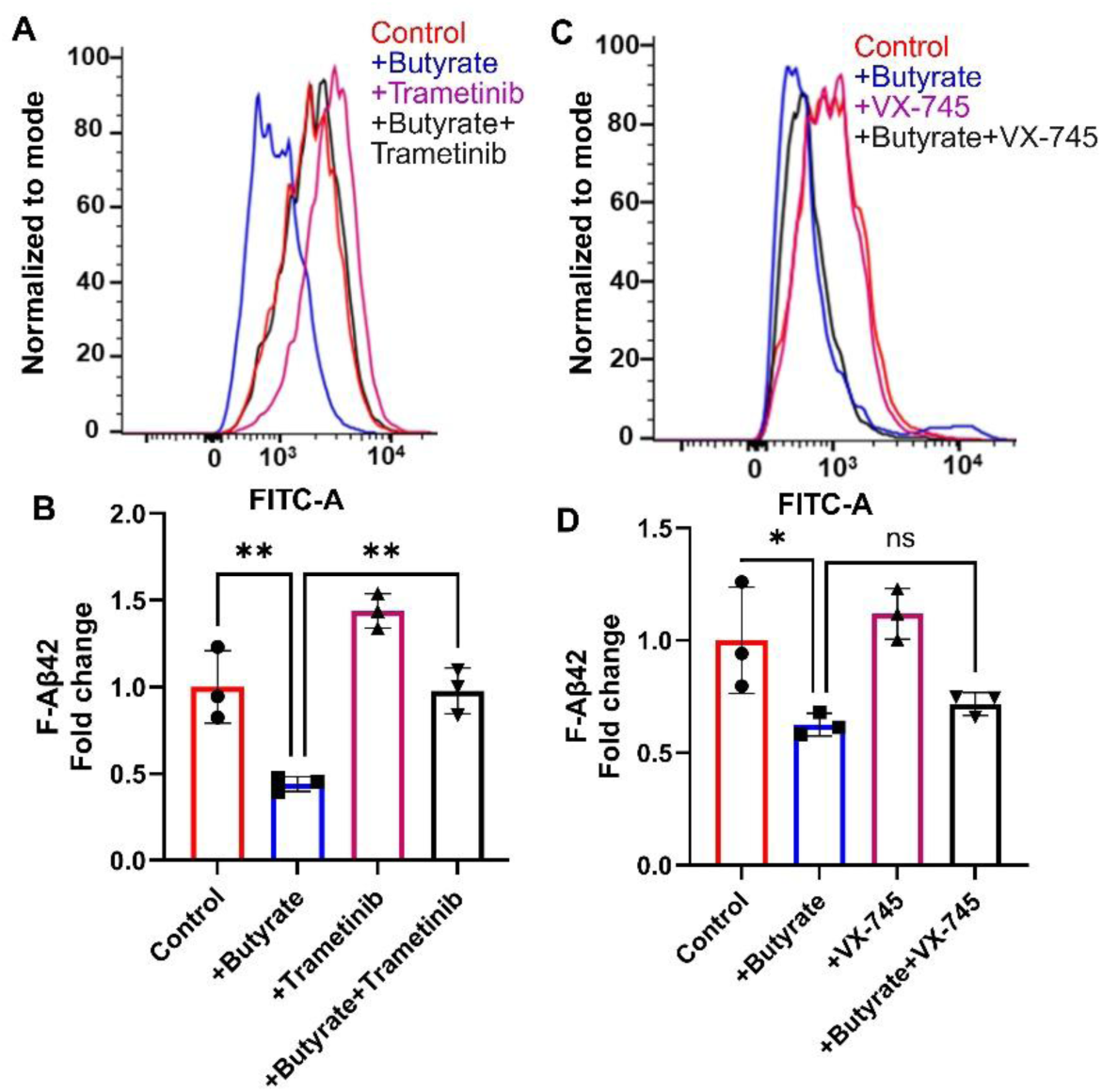
Effect of MEK and p38 inhibitors on butyrate-mediated reduction of F-Aβ42 uptake in polarized hCMEC/D3 monolayers: Cells were incubated for 6 h under the following conditions: control (no treatment), butyrate alone (10 nM), butyrate + inhibitor, or inhibitor alone. In the last 30 minutes of each treatment, all groups received F-Aβ42 (1 μM) and insulin (50 nM). A & B. Butyrate reduced F-Aβ42 accumulation compared to control. Butyrate co-incubation with 0.5 µM trametinib (MEK inhibitor) increased F-Aβ42 accumulation compared to the butyrate alone group, as shown in the histogram of intracellular MFI (A) and bar chart (B) showing fold change of median fluorescence ± standard deviation. C & D. In another experiment, butyrate decreased F-Aβ42 accumulation compared to control, and co-incubation with 10 nM VX-745 (p38 inhibitor) did not alter F-Aβ42 accumulation, as shown in the histogram of intracellular fluorescence (C) and bar chart (D) representing fold change of median fluorescence ± standard deviation. One-way ANOVA followed by Bonferroni post-test; *p-value<0.05, **p-value<0.01.

### 4. Effect of butyrate on transporters/proteins at the BBB in the polarized hCMEC/D3 monolayers

Cells were treated with butyrate for 5 h and co-incubated with Aβ42 for one hour. The Aβ42 treatment decreased P-gp expression in hCMEC/D3 cell monolayers compared to the control; however, this change didn’t meet statistical significance. Co-incubation with butyrate significantly reversed the Aβ42-induced reduction in P-gp levels (p < 0.05), restoring them to the same levels as those in the control (Figure 5A & B). In contrast, Aβ42 treatment significantly increased RAGE expression by 2.4-fold (p<0.05). Butyrate co-incubation reversed the Aβ42-induced increase in RAGE expression, although this effect was not significant (Figure 5A & C).

**Figure 5:**
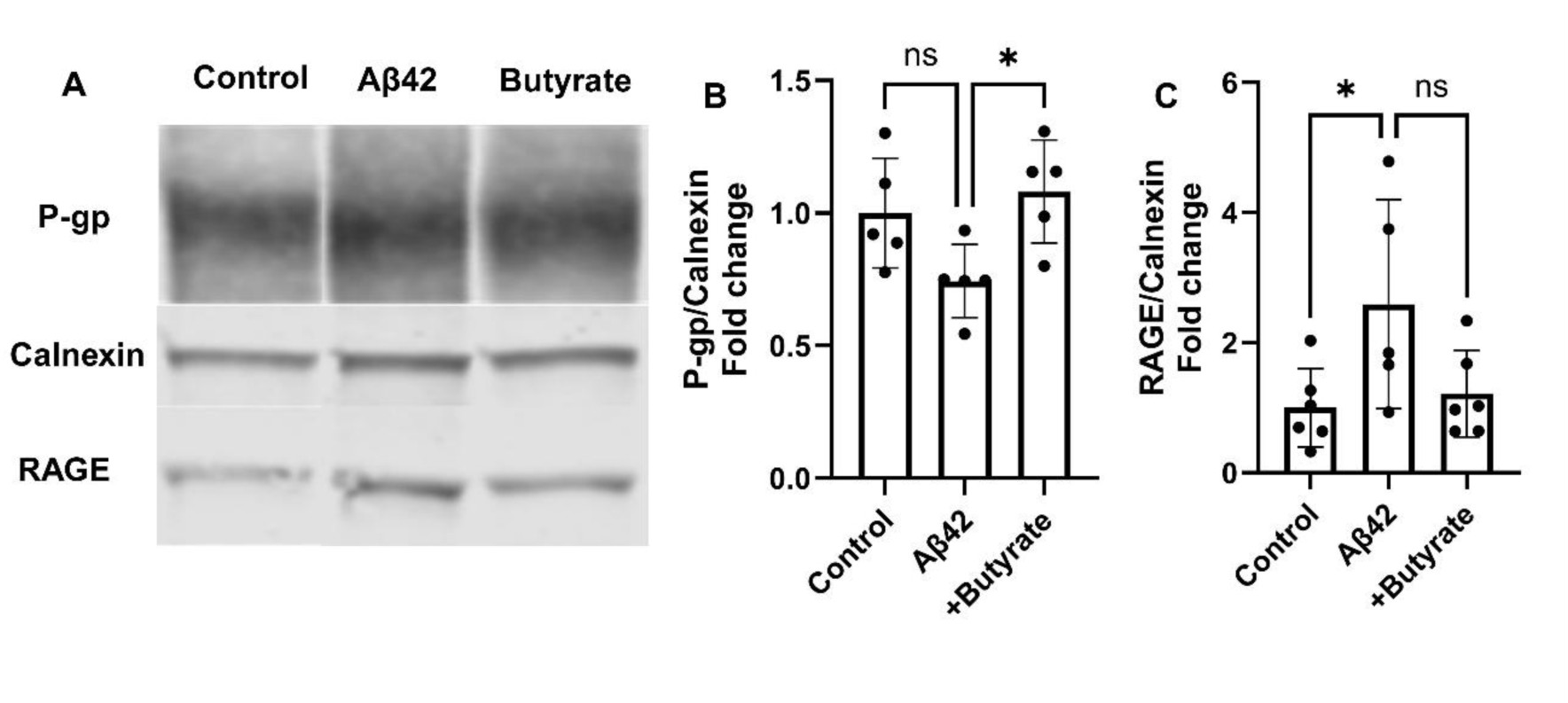
Butyrate effect on the expression of transporters and receptors in polarized hCMEC/D3 monolayers: Cells were treated with 1 μM Aβ42 alone or after butyrate pre-treatment for six hours, with Aβ42 added for the last one hour of incubation. **A.** Immunoblots show that Aβ42 has no significant impact on P-gp expression compared to control; however, butyrate pre-treatment significantly increased P-gp expression compared to the Aβ42 control. On the other hand, Aβ42 significantly increased RAGE expression, but butyrate pre-treatment had no significant impact. Quantification of P-gp (B) and RAGE (C) blots normalized to the expression of calnexin (loading control) was presented as fold change. Mean ± standard deviation; One-way ANOVA followed by Bonferroni post-test; *p-value<0.05.

### 5. Effect of butyrate on Claudin 5 and TIMP-2 levels

The polarized hCMEC/D3 monolayers grown on coverslip dishes were treated with Aβ42 for 1 h, with or without a 6 h butyrate pretreatment. The claudin-5 expression increased upon butyrate pre-treatment compared to the Aβ42 alone group (Figure 6A). The butyrate effect was further investigated in vivo by colonizing germ-free mouse guts with butyrate-producing or KO bacteria. In these mice, we investigated the changes driven by butyrate on various receptors and transporters that mediate Aβ trafficking in and out of the brain. The TIMP-2 levels were measured in the brain cortex of germ-free mice colonized with butyrate KO or butyrate-producing bacteria. The TIMP-2 levels are higher (p<0.01, paired t-test) in the brains of butyrate-producing mice compared to the butyrate KO control (Figure 6B and C).

**Figure 6:**
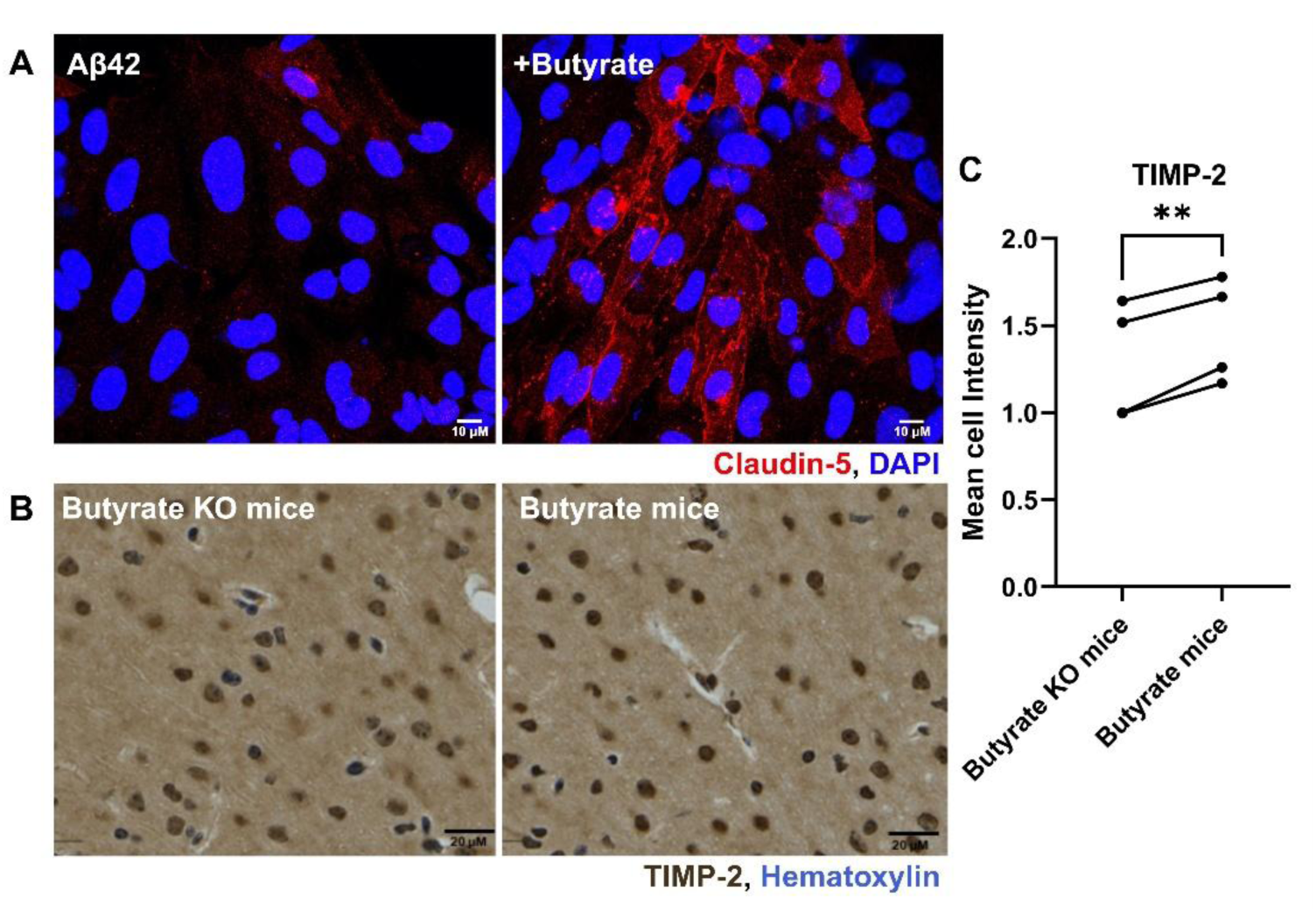
Butyrate effect on claudin-5 and TIMP-2: The polarized hCMEC/D3 monolayers were pre-incubated with butyrate for 6 h, and 1 μM Aβ42 was added for the last hour of incubation. A. Confocal micrographs of the butyrate and Aβ42 co-incubated group show an increase in immunofluorescence pertaining to AF647-tagged claudin-5 compared to the Aβ42 control. TIMP-2 expression was visualized in the brains of butyrate-producing and butyrate-KO bacteria colonized mice through IHC. B. TIMP-2 expression was significantly higher in the brains of butyrate-producing mice compared to the butyrate KO control. The quantification is shown in C as the mean DAB cell pixel intensity ± standard deviation. Paired Student’s t-test, **p-value<0.01.

## Discussion

It is widely recognized that the BBB plays a vital role in clearing Aβ peptides from the brain. In AD, BBB dysfunction is observed, resulting in the disruption of Aβ clearance and leading to increased brain accumulation, which potentially worsens cognitive decline. Hence, it is crucial to investigate treatments that can restore BBB function and eliminate Aβ peptides, maintain homeostasis.

The current study has demonstrated that 10 nM butyrate significantly reduces the F-Aβ42 uptake by BBB cell culture models (hCMEC/D3 and BBME monolayers) (Figure 1). The butyrate showed a more pronounced effect on the endothelial accumulation of toxic amyloidogenic F-Aβ42 compared to F-Aβ40. Similarly, butyrate also reduced the Aβ-induced neurotoxicity in a neuronal cell culture model^33^. The butyrate effect was particularly evident in the shorter time frames (2 h and 6 h) following incubation; the absence of detectable butyrate effect 24 h after incubation could be due to the metabolic utilization of butyrate as an energy substrate by the endothelial cells. This phenomenon is consistent with previous observations reported in colonocytes and hepatocytes^34,35^. Similar to the in vitro results, butyrate-fed AD transgenic mice (5xFAD) also exhibited reduced plaque deposition and improved behavioral assessments compared to the control^36^. These findings further strengthen the fact that butyrate treatment mitigates Aβ-induced toxicity, as reported in various models^37–40^. Alternatively, claims have been made that short-chain fatty acids derived from the microbiota promote Aβ plaque deposition in germ-free AD mice, although the effects of individual metabolites were not explored^41^. Hence, to understand how butyrate reduces Aβ uptake, it is critical to investigate the impact of butyrate on the BBB endothelium and its underlying mechanisms.

### Effect of butyrate on insulin signaling at the BBB

Metabolic syndrome, characterized by insulin resistance and disrupted insulin signaling, is one of the major risk factors for Alzheimer’s disease ^42–45^. Restoring the insulin signaling in cerebral vasculature, particularly at the BBB endothelium, is critical for developing neuroprotective strategies. Butyrate enhances insulin sensitivity by phosphorylating Insulin Receptor Substrate-1 and AKT, through histone acetylation.^46^ Furthermore, butyrate has been shown to modulate ERK phosphorylation in hypoxic-ischemic brain injury^47^. Based on these reports, we hypothesized that butyrate could enhance insulin signaling in BBB endothelial cells. We focused on the two major arms of the insulin signaling pathway: the PI3K/AKT pathway, which regulates energy metabolism, and ERK/MAPK pathway, which controls cell proliferation, growth, and differentiation^48,49^. To accurately assess changes in insulin signaling, despite the inherent limitations of Western blot sensitivity, cells were stimulated with insulin with and without butyrate. Western blot analysis of BBB endothelial cell lysates showed that treatment with both butyrate and insulin significantly increased phosphorylation of AKT and ERK compared with the insulin-alone group (Figure 2). These studies demonstrate that butyrate potentiates insulin signaling in the BBB endothelium. The activation of AKT and ERK by butyrate may have functional consequences for BBB physiology and its regulation of cellular transport and endocytosis^50^. Therefore, we conducted additional studies to investigate the critical link between this butyrate-mediated activation of AKT and ERK and its potential role in regulating F-Aβ42 uptake by BBB endothelial cells.

### Role of butyrate-activated insulin signaling pathways in Aβ42 trafficking at the BBB

#### The PI3K/AKT Pathway and F-Aβ42 uptake

Co-incubation of BBB endothelial cells with butyrate and MK2206, a selective AKT inhibitor, significantly increased F-Aβ42 accumulation compared to the cells treated with butyrate alone (Figure 3). This key finding indicates that the butyrate-mediated decrease in F-Aβ42 uptake was mediated through the phosphorylation of AKT (Ser473) and activation (Figure 3). Aβ42 is known to impair AKT signaling, thereby suppressing protective mechanisms and promoting Aβ toxicity as well as neuronal loss^51^. Our data suggest that butyrate actively reverses the Aβ-mediated suppression of AKT at the BBB. We also investigated the role of mTORC1, a downstream node of the AKT pathway, in F-Aβ42 uptake by treating hCMEC/D3 cell monolayers with rapamycin, an mTORC1 inhibitor. Co-incubation of rapamycin, with butyrate did not affect F-Aβ42 uptake compared to the butyrate treatment alone (Figure 3). This result shows that the butyrate-mediated reduction of F-Aβ42 accumulation is independent of mTORC1 phosphorylation, although AKT is involved. Moreover, increased mTOR activity has been cited in literature as potentially worsening Aβ pathology, possibly due to other pathways activated by amino acids, cholesterol trafficking, and adenosine 5’-monophosphate-activated protein kinase^52,53^.

#### The ERK/MAPK pathway and F-Aβ42 uptake

Next, we examined the role of the major MAPK kinases (ERK and p38) on F-Aβ42 uptake by the BBB endothelial cells. The role of butyrate-mediated activation of major MAPK kinases-ERK and p38 in F-Aβ42 uptake was investigated by treating cells with specific inhibitors. We found that the co-incubation of Trametinib, an ERK inhibitor, and butyrate increased F-Aβ42 uptake compared to the butyrate-alone group (Figure 4), which indicates that butyrate-driven ERK phosphorylation actively decreases F-Aβ42 uptake in the hCMEC/D3 cell monolayers. This finding contrasts with some literature reports, where β-amyloid precursor protein-induced ERK phosphorylation is linked to impaired signaling and exacerbated Aβ pathology in microglia^54^. This contradiction may be due to key differences, including the extent of ERK phosphorylation achieved or differences in cell types (endothelial cells vs. microglia). Furthermore, co-incubation with another MAPK kinase-p38 inhibitor (VX-745) didn’t affect F-Aβ42 uptake compared to the butyrate alone group (Figure 4), ruling out P38 as a mediator of the butyrate effect. Taken together, these findings provide a strong mechanistic basis for concluding that butyrate modulates Aβ uptake in the BBB endothelium via the AKT and ERK pathways, rather than mTORC1 and p38 pathways. Therefore, it is highly probable that mechanisms regulating Aβ trafficking are disrupted in insulin resistance, leading to toxic Aβ accumulation within endothelial cells.

### Butyrate modulates key Aβ receptors and transporters and restores BBB integrity

We then examined butyrate’s effect on the expression of major Aβ transporters and receptors in polarized hCMEC/D3 cell monolayers. The effect of butyrate on transporters and receptors that impact Aβ transport was investigated in polarized hCMEC/D3 cell monolayers. LRP1 and P-gp mediate Aβ clearance from the brain, whereas RAGE mediates the influx^55,56^. In AD human brains, increased Aβ deposition is directly correlated with RAGE and inversely with P-gp and LRP expression; RAGE expression was increased, whereas P-gp and LRP expression has been reported to decrease^6,57,58^. We observed similar trends in the hCMEC/D3 cell culture model. Aβ42 treatment led to a reduction in P-gp levels (though not statistically significant), and this effect was reversed upon co-incubation with butyrate, restoring P-gp to control levels (Figure 5). RAGE expression was significantly increased with Aβ42 treatment compared to control. Butyrate co-incubation decreased RAGE expression compared to Aβ42 alone group, though this did not meet statistical significance. This lack of significance could be attributed to the relatively shorter duration of the incubation period.

### Butyrate protects tight junctions and barrier integrity

Tight junctions and basement membrane proteins interact to maintain the integrity of the BBB. We have demonstrated that butyrate aids in preserving BBB integrity by modulating junctional protein expression and regulating the composition of basement membrane proteins.

The hCMEC/D3 cell monolayers co-incubated with butyrate upregulated claudin-5, a tight junction protein, compared to the Aβ42-treated group. It demonstrates butyrate’s efficacy in alleviating the Aβ42’s detrimental impact on junctional integrity (Figure 6). This protective effect may be linked to its influence on the composition of the basement membrane. Butyrate increases the TIMP-2 levels in germ-free mice colonized with butyrate-producing bacteria. The Aβ peptides are known to increase the production of matrix metalloproteinases (MMP), which degrade extracellular matrix components, thereby disrupting tight junction proteins such as claudin-5^59^. Since TIMP-2 is a potent inhibitor of MMPs, this increase is likely a key mechanism by which butyrate restores claudin-5 expression. Furthermore, TIMP2 has been shown to activate AKT upon interacting with membrane-type MMP in hCMEC/D3 monolayers^60^. Given our earlier finding that butyrate activates AKT, the pathway provides a potential link that TIMP-2 drives AKT activation, which in turn upregulates P-gp, thereby enhancing Aβ42 efflux.

It is important to note that a complementary study conducted in germ-free mice colonized with butyrate-producing bacteria showed no alteration in the key components of BBB trafficking, including caveolin-1, vesicular transcytosis inhibitor-MFSD2A, and transporters implicated in Aβ42 uptake, such as P-gp (Supplementary figure). This suggests that the P-gp and RAGE modulatory effects of butyrate are most **prominent in the presence of Aβ pathology**, where butyrate acts to restore BBB dysfunction upon Aβ exposure rather than altering baseline transporter expression in a healthy state.

## Conclusions

In summary, this study provides compelling mechanistic evidence that butyrate decreases pathological Aβ42 uptake via insulin signaling nodes, specifically, AKT and ERK. Activation of this signaling initiates the reversal of BBB dysfunction by increasing the expression of key efflux transporter-P-gp, which promotes Aβ42 efflux, and by restoring claudin-5, a tight junction protein that’s critical for BBB integrity. Butyrate also upregulates TIMP-2, an inhibitor of extracellular matrix-degrading enzymes, in germ-free mice colonized with butyrate-producing bacteria, further strengthening its protective role. Taken together, this mechanistic information highlights the therapeutic potential of butyrate in restoring BBB function and decreasing cerebrovascular amyloid accumulation, meeting criteria for effective treatment. Our future studies will focus on validating the mechanisms identified in this *in vitro* model within relevant animal models.

## Acknowledgements

**Funding:** This research was funded, in part, by the National Institute of Health/National Institute of Neurological Disorders and Stroke R01NS125437 (K.K.K.) and National Institute of Health DK114007 (P.K.C.). Imaging work was supported by the resources and staff at the University of Minnesota University Imaging Centers (UIC)-SCR_020997. We are grateful for a grant-in-aid (#324930) from the University of Minnesota for the purchase of the LI-COR Odyssey CLx Infrared imaging system that supported western blot imaging.

## Author Contributions

This study was conceptualized and designed by K.K.K., V.V., Z.W., P.C.K.; Data acquisition was performed by V.V.; Data analysis was performed by V.V., Z.W. The Manuscript was written by V.V., which was revised and approved by K.K.K., P.C.K., and Z.W.; All authors read and agreed to the final version of the manuscript.

## Abbreviations

AD: Alzheimer’s disease
Aꞵ: Amyloid beta
AF647: Alexa fluor 647
BBB: Blood-Brain Barrier
BBME: Bovine Brain Microvascular Endothelial
DAPI: 4’,6-diamidino-2-phenylindole
DMEM: Dulbecco’s Modified Eagle’s Medium
ERK: Extracellular Signal-Regulated Kinase
F: Fluorescein Isothiocyanate
HFIP: 1,1,1,3,3,3, - hexafluoro-2-propanol
HDAC: Histone deacetylase
hCMEC/D3: Human cerebral microvascular endothelial cells
LRP1: Lipoprotein Receptor-related Protein 1
MAPK: Mitogen-Activated Protein Kinase
MFI: Median Fluorescence Intensity
mTOR: Mammalian Target of Rapamycin
MMP: Matrix Metalloproteinases
PFA: Paraformaldehyde
P-gp: P-glycoprotein
RAGE: Receptor for Advanced Glycation End products
TIMP-2: Tissue Inhibitor of Metalloproteinases-2
TBS: Tris-buffered saline
TBST: Tris-buffered saline containing Tween 20

