## Supplementary figure for "Butyrate regulates the blood-brain barrier transport and intra-endothelial accumulation of Alzheimer’s disease Amyloid-beta peptides"

### **Supplementary information**

**Methods:**

**Immunohistochemistry (IHC)**

The mice’s brains were formalin-fixed and embedded in paraffin, and then sectioned into 5-µm-thick slices on a slide. Sections were processed for IHC as previously published^36^. Brain tissue was stained for tissue inhibitors of metalloproteinases (P-gp, 1:500 dilution, #ab170904 Abcam, Cambridge, MA, caveolin-1, 1:250 dilution, #3238, MFSD2A, 1:500 dilution, #80302 from Cell Signaling Technology, MA) with 3,3′-Diaminobenzidine (DAB) chromogen. Images were acquired using a Zeiss Axio Scan.Z1 microscope at 20X magnification. Images were analyzed in QuPath-0.5.1 where germ-free mouse brains were compared to butyrate-producing microbiome colonized mouse brains. Mean DAB pixel intensity from both groups was obtained by averaging over 100 cell-specific regions of interest in the brain cortex, followed by pair-wise comparison.

**Results:**

P-gp, caveolin-1 and MFSD2A expression didn’t change significantly between germ-free and butyrate-producing microbiome colonized mice brains. Representative images were shown in Figure S1.

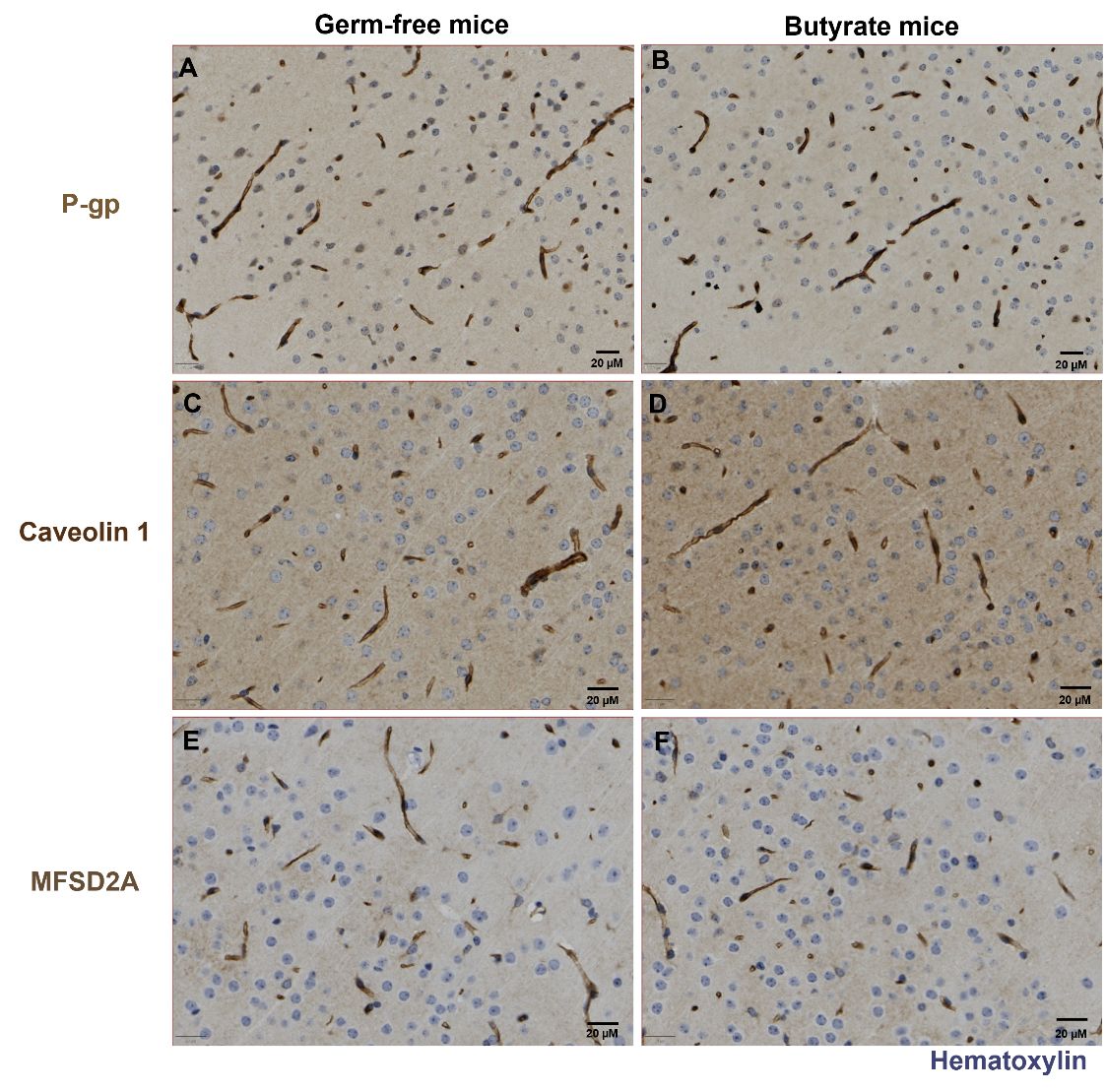

**Figure S1:** Germ-free and butyrate-producing microbiome colonized mice brains were probed for various targets involved in Aβ42 trafficking regulation. Expression levels of P-gp (A, B), caveolin-1 (C, D), and MFSD2A (E, F) are not varied between the two comparative groups.
